# Exceptional aggregation propensity of amino acids in polyglutamine amino-acid-homopolymer

**DOI:** 10.1101/2020.07.09.194753

**Authors:** Rahul Mishra, Ashwani K. Thakur

**Affiliations:** Department of Biological Sciences and Bioengineering, Indian Institute of Technology, Kanpur, India-208016

**Author notes:** To whom correspondence should be addressed: Prof. Ashwani Kumar Thakur.

**Keywords:** Polyglutamine, Aggregation, β-sheet-propensity

## Abstract

Similar aggregation and β-sheet propensity of amino acids in globular proteins and amyloids, suggests comparable principles of their formation. Here we show that during the process of aggregation into amyloid-like fibers, these rules are not the same in an amino-acid-homopolymer (AAHP) polyglutamine (PolyGln). An aggregation kinetic analysis on nine-point mutants of a forty-six long PolyGln peptide was carried in physiological conditions. At the dynamic equilibrium state of aggregation, critical-concentration derived free-energy differences, signifying aggregation propensity of incorporated amino acids were obtained. None of the obtained propensities correlated with existing conventional aggregation and β-sheet propensities of the amino acids in proteins and amyloids. Further, the differential aggregation behavior of all the peptides only correlated with van der Waals volume of the incorporated amino acid and not with any other physicochemical characteristic of amino acids. The new rules obtained from PolyGln AAHP provide an opportunity to explore physiological relevance of a mutation within AAHP in human proteome. Additionally, this study opens up new avenues for protein model design exploring folding and aggregation behavior of other amino-acid-homopolymer (AAHP) existing in the human proteome.

**Significance:** Mutational analysis within PolyGln sequences adds to the knowledge of unique aggregation propensities of amino acids within PolyGln AAHP. This study highlights the importance of van der Waals volume in dictating stability-instability of an aggregation fold and in turn aggregation kinetics and thermodynamic stability of aggregates. The analysis signifies the role of Gln-Gln interlocking system within PolyGln folding motif and extent of disruption caused by van der Waals volume of an amino acid. The results can be taken as a starting point to evaluate the possible impact of amino acid insertions in PolyGln stretches of other proteins. It also opens opportunities to study the structural and functional relationship of other AAHPS for their unique folding and aggregation behavior. Learning outcome can be utilized as a bottom–up approach to design amyloid biomaterial with different strengths for biomedical applications.

---

Polyglutamines (PolyGlns) represent a unique class of amino acid homopolymer (AAHP) in cellular context(1). Expansion beyond a threshold length spontaneously aggregate them and the aggregation is associated with at least ten neurodegenerative diseases including Huntington’s disease(2). Structurally, PolyGln aggregates exhibit striking similarities with amyloid fibers generated from amyloidic prone and globular proteins. Electron microscopy (EM) examination of PolyGln aggregates displays morphological structures similar to amyloid fibrils (3, 4). Biochemical investigation of PolyGln aggregates demonstrates thioflavin T (ThT) binding and shifting of the fluorescence spectrum similar to regular amyloid aggregates (4). PolyGln aggregates also showed binding towards amyloid-specific monoclonal antibodies (4). PolyGln aggregation pathway follows nucleation aggregation kinetics model (4-6), analogous to other amyloidogenic peptides (7).

Notably, protein aggregation is a generic nature of any polypeptide chain and it seems that similar sets of rules govern the formation of β-sheet rich amyloid aggregates (8). The tendency of proteins and peptides to aggregate is linked with decrease in overall net charge, increase in hydrophobicity and high β-sheet propensity of amino acids present in them (9). Such rules hold true for amyloid aggregates generated from both amyloid and globular proteins. These rules are deciphered using mutational studies of amyloid β peptide (Aβ) (10-12), muscle acyl phosphatase protein (13, 14) and α-synuclein (15) protein model systems. According to these studies, bulky hydrophobic amino acids: Ile, Val, Leu along with aromatic residues: Trp, Phe and Tyr have very high aggregation propensity. Charge residues universally disfavor the aggregation and amyloid formation. Interestingly, the aggregation propensity scale of the above mentioned amino acids were also found to match with their β-sheet propensity scale (12) (16), (17). Therefore, it appears that amino acids use similar principles towards formation of β-sheet rich amyloid aggregates and β-sheet in globular proteins.

Contrary to the striking similarities between amyloid aggregates and PolyGln amyloid like fibrillar structures, it is still not clear how amino acids with high hydrophobicity, lower net charge and high intrinsic β-sheet propensity will influence the PolyGln aggregation behaviour. One of the main difficulties in conducting such mutational analysis in the PolyGln context is the absence of a proper PolyGln protein/peptide model system which can provide structurally interpretable results. The main drawback of conducting a mutational analysis using a PolyGln disease protein model system is that the PolyGln aggregation behavior can be easily influenced by the amino acids flanking the PolyGln stretch (18). Thus interpretation of amino acids’ physicochemical factors contributing towards promoting or restraining the PolyGln aggregation becomes challenging (2, 19). Similarly, choosing an aggregation prone PolyGln peptide of pathological length for mutational analysis will have a tendency to overpass and adjust a mutation due to their polar zipper character (19). This may pose another challenge to evaluate the exact impact of an amino acid interruption on the PolyGln aggregation kinetics.

To overcome these PolyGln protein/peptide model related limitations, we choose an aggregation prone PolyGln peptide model (PepQ; 46 residue length, Figure 1) interrupted with proline-glycine (Pro-Gly) at regular intervals(19). Introduction of (Pro-Gly), a β-turn inducing pair after every nine Glns (Q_9_) are expected to avoid slippage of PolyGln sequences over each other and thereby maintaining the antiparallel β-sheet folding motif(19). Choosing a PolyGln model system with an antiparallel β-sheet folding motif is important as it also provides a chance to understand our results in the context of recently determined antiparallel β-sheet rich PolyGln aggregate core structure (20). However, before using the PolyGln peptide model for mutation experiments, its efficiency to reflect the PolyGln aggregation kinetics, according to the nature of amino acid inserted was tested.

**Figure 1.**
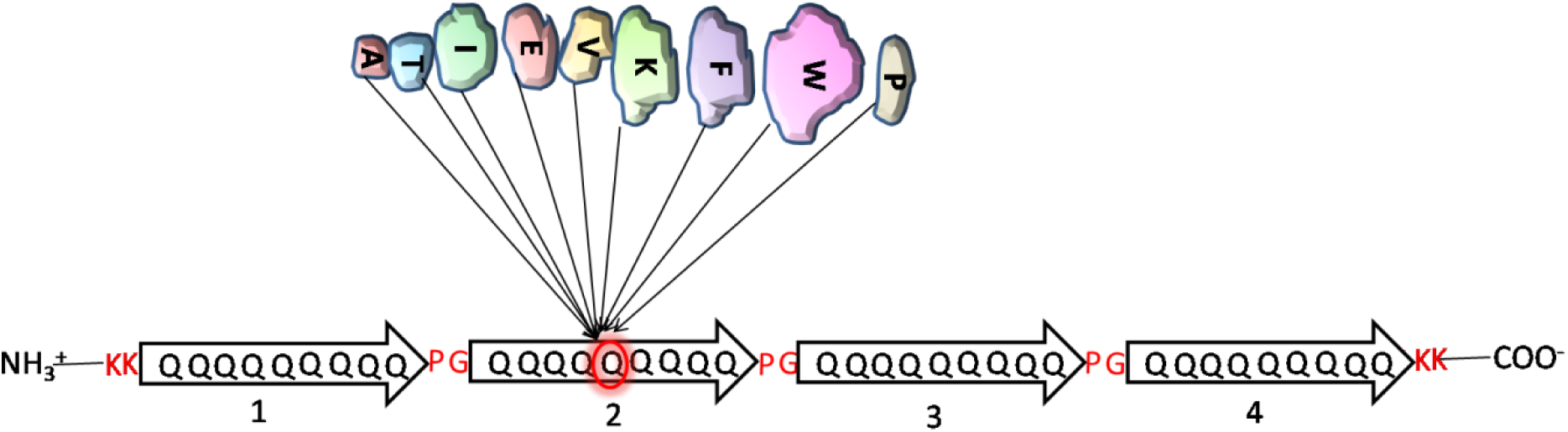
Schematic representation of a forty-six residue PolyGln peptide model system, where Gln at the 18^th^ position is substituted with nine different amino acids (Table 1).

To test the PolyGln model efficiency, initially Pro (β-strand breaking), Lys (positively charge) and Glu (negatively charge) amino acids were substituted for Gln at 18^th^ position in the PepQ peptide (refer Table 1 for sequence details). Depending on the physicochemical nature of these residues, aggregation kinetics of PepP, PepK and PepE were compared against PepQ peptide. In PepP, insertion of Pro does not allow PepP to nucleate and aggregate (19). This result matches with the Pro amino acid capability to block amyloid formation as it is strongly disfavored in the interior strands of β-sheet. Similarly, as charge residues presence also resists protein aggregation, PepK and PepE spontaneous aggregation kinetics was found to be intermediate of PepQ and PepP (21). Additionally, when a salt-bridge was allowed to form between Lys and Glu of PepK and PepE peptides resulting in neutralization of their charges, nucleation pathways of both peptides changed to form nanospheres as compared to fibers (21). Aggregation kinetics of PepP, PepK and PepE clearly demonstrate that mutational analysis of the aggregation behavior of PolyGln, with different amino acids at 18^th^ position can be analyzed within two extremes: aggregation (Gln^18th^, PepQ) and non-aggregation (Pro^18th^, PepP).

**Table 1.**
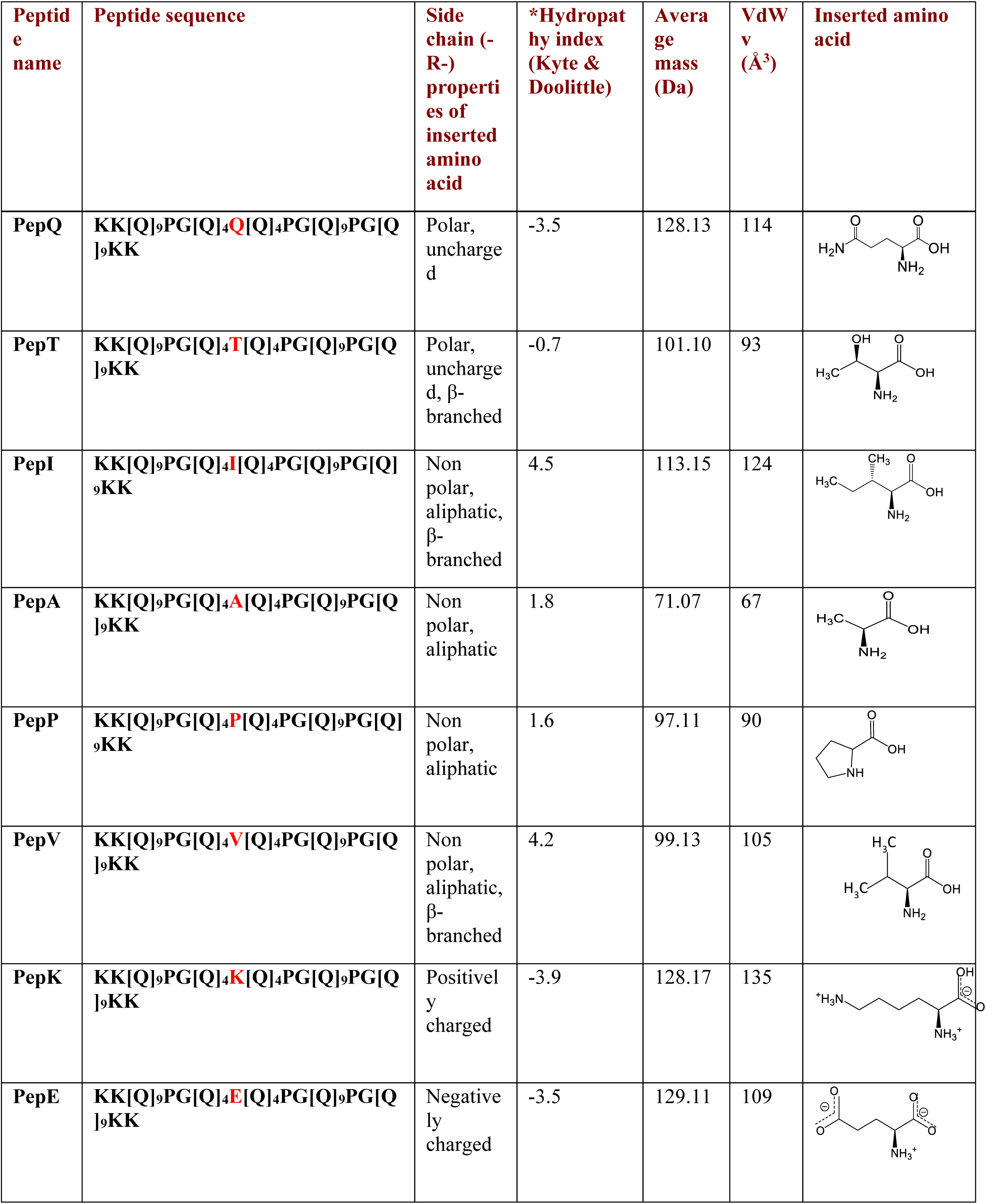
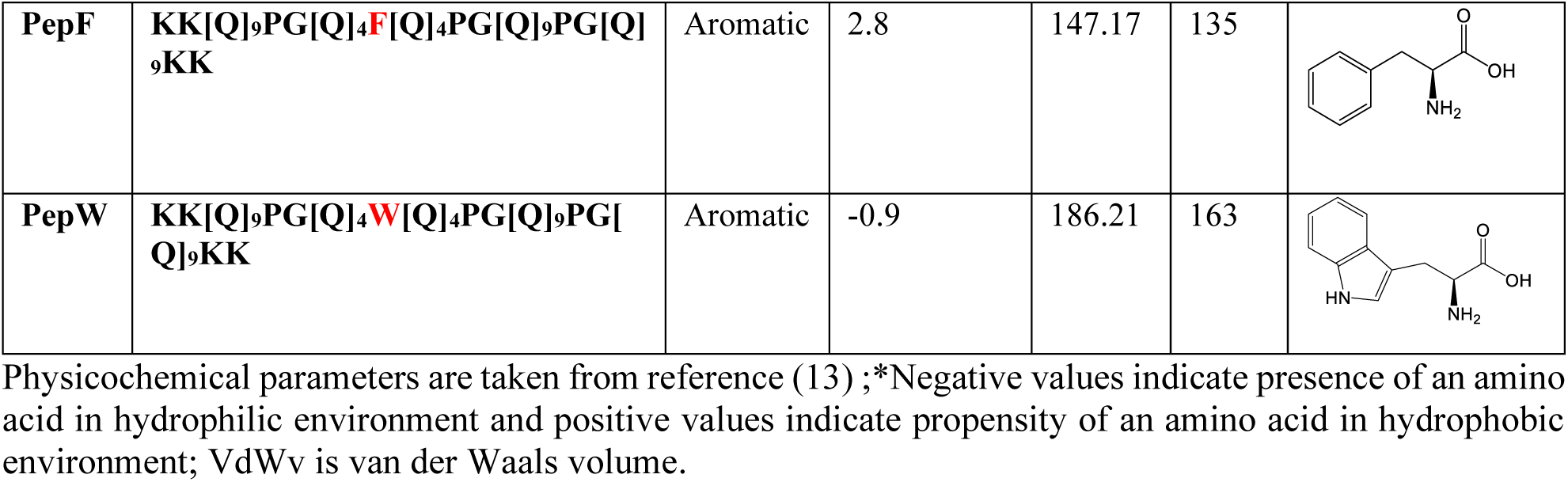
Physicochemical properties of amino acids inserted in PolyGln peptides.

| Peptide name | Peptide sequence | Side chain (-R-) properties of inserted amino acid | *Hydropathy index (Kyte & Doolittle) | Average mass (Da) | VdW v (Å<sup>3</sup>) | Inserted amino acid |
| --- | --- | --- | --- | --- | --- | --- |
| PepQ | KK[Q]<sub>9</sub>PG[Q]<sub>4</sub>**Q**[Q]<sub>4</sub>PG[Q]<sub>9</sub>PG[Q]<sub>9</sub>KK | Polar, uncharged | -3.5 | 128.13 | 114 |  |
| PepT | KK[Q]<sub>9</sub>PG[Q]<sub>4</sub>**T**[Q]<sub>4</sub>PG[Q]<sub>9</sub>PG[Q]<sub>9</sub>KK | Polar, uncharged, β-branched | -0.7 | 101.10 | 93 |  |
| PepI | KK[Q]<sub>9</sub>PG[Q]<sub>4</sub>**I**[Q]<sub>4</sub>PG[Q]<sub>9</sub>PG[Q]<sub>9</sub>KK | Non polar, aliphatic, β-branched | 4.5 | 113.15 | 124 |  |
| PepA | KK[Q]<sub>9</sub>PG[Q]<sub>4</sub>**A**[Q]<sub>4</sub>PG[Q]<sub>9</sub>PG[Q]<sub>9</sub>KK | Non polar, aliphatic | 1.8 | 71.07 | 67 |  |
| PepP | KK[Q]<sub>9</sub>PG[Q]<sub>4</sub>**P**[Q]<sub>4</sub>PG[Q]<sub>9</sub>PG[Q]<sub>9</sub>KK | Non polar, aliphatic | 1.6 | 97.11 | 90 |  |
| PepV | KK[Q]<sub>9</sub>PG[Q]<sub>4</sub>**V**[Q]<sub>4</sub>PG[Q]<sub>9</sub>PG[Q]<sub>9</sub>KK | Non polar, aliphatic, β-branched | 4.2 | 99.13 | 105 |  |
| PepK | KK[Q]<sub>9</sub>PG[Q]<sub>4</sub>**K**[Q]<sub>4</sub>PG[Q]<sub>9</sub>PG[Q]<sub>9</sub>KK | Positively charged | -3.9 | 128.17 | 135 |  |
| PepE | KK[Q]<sub>9</sub>PG[Q]<sub>4</sub>**E**[Q]<sub>4</sub>PG[Q]<sub>9</sub>PG[Q]<sub>9</sub>KK | Negatively charged | -3.5 | 129.11 | 109 |  |
| **PepF** | **KK[Q]<sub>9</sub>PG[Q]<sub>4</sub>F[Q]<sub>4</sub>PG[Q]<sub>9</sub>PG[Q]<sub>9</sub>KK** | Aromatic | 2.8 | 147.17 | 135 |  |
| **PepW** | **KK[Q]<sub>9</sub>PG[Q]<sub>4</sub>W[Q]<sub>4</sub>PG[Q]<sub>9</sub>PG[Q]<sub>9</sub>KK** | Aromatic | -0.9 | 186.21 | 163 |  |

Physicochemical parameters are taken from reference (13) ;\*Negative values indicate presence of an amino acid in hydrophilic environment and positive values indicate propensity of an amino acid in hydrophobic environment; VdWv is van der Waals volume.

In this paper, besides PepQ and PepP, eight PolyGln peptides where Gln at 18^th^ position was substituted with different amino acids (Ala, Ile, Thr, Val, Lys, Glu, Phe, Trp) and studied for aggregation reaction kinetics. Using this mutational data, we answered two main questions: 1) will the aggregation propensity rules of amino acids obtained from PolyGln AAHP match with the propensity scale from other amyloid proteins, 2) which physicochemical factor of amino acids will govern the PolyGln aggregation.

Results presented here revealed that the aggregation propensity rules obtained from PolyGln AAHP showed no correlation with the scale from other amyloid and globular proteins. Size of the amino acids (van der Waals volume) incorporated in the PolyGln peptide model, controls the hierarchical levels of PolyGln aggregation, critical concentration, thermodynamic stability of aggregates and the core structure.

## Results

### Spontaneous aggregation kinetics

Purified peptides after disaggregation and solubilization were subjected to spontaneous aggregation reaction at 37 °C in PBS, pH 7.2. An aliquot of an ongoing reaction was ultracentrifuged and supernatant was used to measure the decrease in monomer concentration with time by RP-HPLC based assay (22) (Figure S1). The concentrations of all the peptides were measured with the help of a standard curve made for each peptide (Figure S2). Aggregation behavior obtained for different mutants varied with respect to PepQ (Figure 2A). This peptide aggregates within 200 hr. PepP as shown earlier did not aggregate within the concentration range used in this study (19). Depending on the overall aggregation rate comparison (Figure 2B, Table S1), of the peptides: PepA, PepT and PepI peptides were considered fast as their aggregation behavior was found close to PepQ. PepE & PepV were found to be moderate in aggregation. PepK, PepF and PepW peptides were slow, while PepP was non-aggregating (Figure 2B).

**Figure 2.**
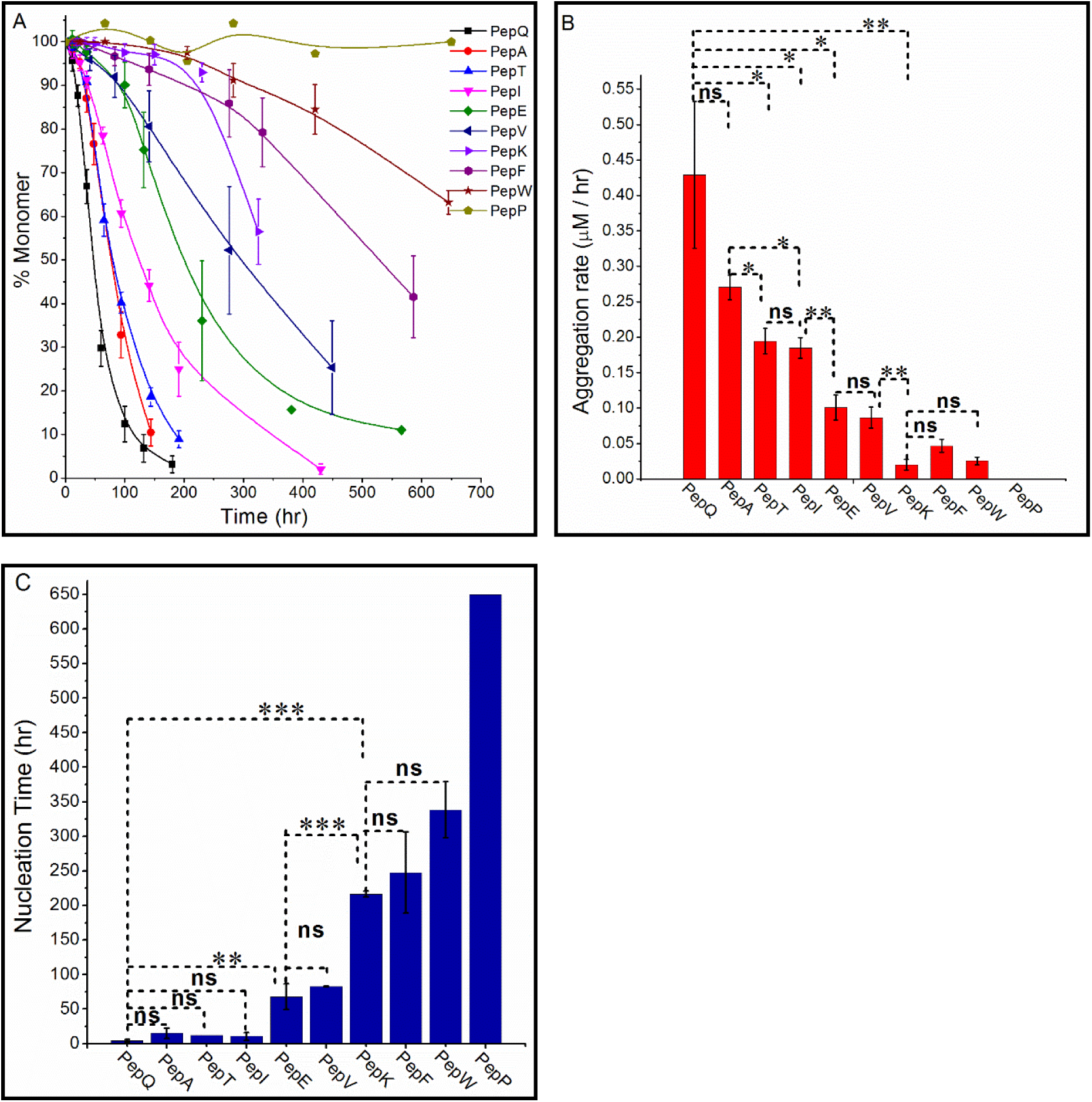
A) Spontaneous aggregation kinetics of different PolyGln mutant peptides monitored at 40 µM concentration through RP-HPLC assay. B) Aggregation rates of all the PolyGln mutant peptides. C) Nucelation time (lag time before aggregation) of PolyGln peptides calculated using RP-HPLC spontaneous aggregation profile. Statistical significance of measurement is represented by asterisks (***, P<0.0001; **, 0.001 < P < 0.01; *, 0.01 < P < 0.05; ns = not significant, P > 0.05). For all the experiments, error bars represent standard error of mean for minimum n = 4, except for PepP where n = 1. As PepP does not aggregate, its nucleation time has no meaning. The time represented here is to show maximum aggregation monitoring duration of PepP.

As compared to all other mutants, PepQ aggregated much faster, with majority of its monomers, exhausting within 200 hr (Figure 2A) and have aggregation rate of 0.43 ± 0.10 μM/hr (Figure 2B). This is followed closely by PepA, PepT and PepI with aggregation rate of 0.27 ± 0.02 μM/hr, 0.19 ± 0.02 μM/hr and 0.18 ± 0.01 μM/hr, respectively (Figure 2B, see Table S1 for aggregation rates of all the mutants). By looking at the differential behavior of aggregation, it appears that size of the substituted amino acid side chain shows its influence on spontaneous aggregation kinetics (elaborated in the discussion section). For example, Ala and Thr side chains are smaller in size as compared to Gln and can be adjusted efficiently in the PolyGln aggregation prone motif and thus aggregates fast (Table 1). Similarly, Lys and Phe in PepK and PepF peptides have larger size, consequently aggregates slowly. Likewise, Trp in PepW has the largest size (Table 1) and sluggishly aggregates. Further, different nucleation times (Figure 2C) obtained suggest that the side chains of the substituted amino acids in these peptides cause constraints to different extents, in the aggregation prone folding motif (23). For example, Ala, Thr and Ile amino acids seem less disruptive as compared to moderate (Glu, Val) and slow aggregators (Lys, Phe and Trp) in comparison to PepQ.

### Free energy of fibril formation through critical concentration (Cr) determination

It is known from the mutational experiments on different amyloid proteins and peptides that substitutions of amino acids influence the thermodynamic stability of amyloid fibers (15, 24-27). Thermodynamic stability can be calculated by using critical concentration (*Cr*) values (Table S2). It is the residual monomer concentration present at the end of an aggregation reaction and represents the threshold concentration of protein/peptide molecules, above which it can aggregate (28). *Cr* values of all the mutants were calculated by spontaneous and seeding aggregation kinetics in the forward direction and by spontaneous dissociation kinetics in the reverse direction (Figure 3). As expected from the operation of a dynamic equilibrium at the reaction endpoint, both directions yielded similar values of critical concentration. The values of all the peptides range from 0.25 ± 0.01 μM to 7.9 ± 0.42 μM (Table S2). These values were then converted to thermodynamic parameter 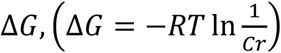, which corresponds to the free energy and thus thermodynamic stability of fibril formation (11, 22). Rapidly aggregating PolyGln sequences gave less *Cr* values and accordingly more negative ΔG values in comparison to slow aggregating PolyGln mutants. These values range from -9.36 ± 0.05 kcal/mol to -7.24 ± 0.07 kcal/mol (Table 3). Fast aggregating mutants PepA (−9.34 ± 0.05 kcal/mol), PepT (−8.93 ± 0.32 kcal/mol) and PepI (−8.98 ± 0.05 kcal/mol) have ΔG values close to the host peptide PepQ (−9.36 ± 0.05 kcal/mol). Similarly, slow aggregating mutants have less negative ΔG values. It means that fibers of fast aggregating mutants are expected to be more stable as compared to slow aggregating mutants (22, 28). The values obtained for the PolyGln mutant peptides were similar to those reported earlier from PolyGln peptides of equivalent length (29).

**Figure 3.**
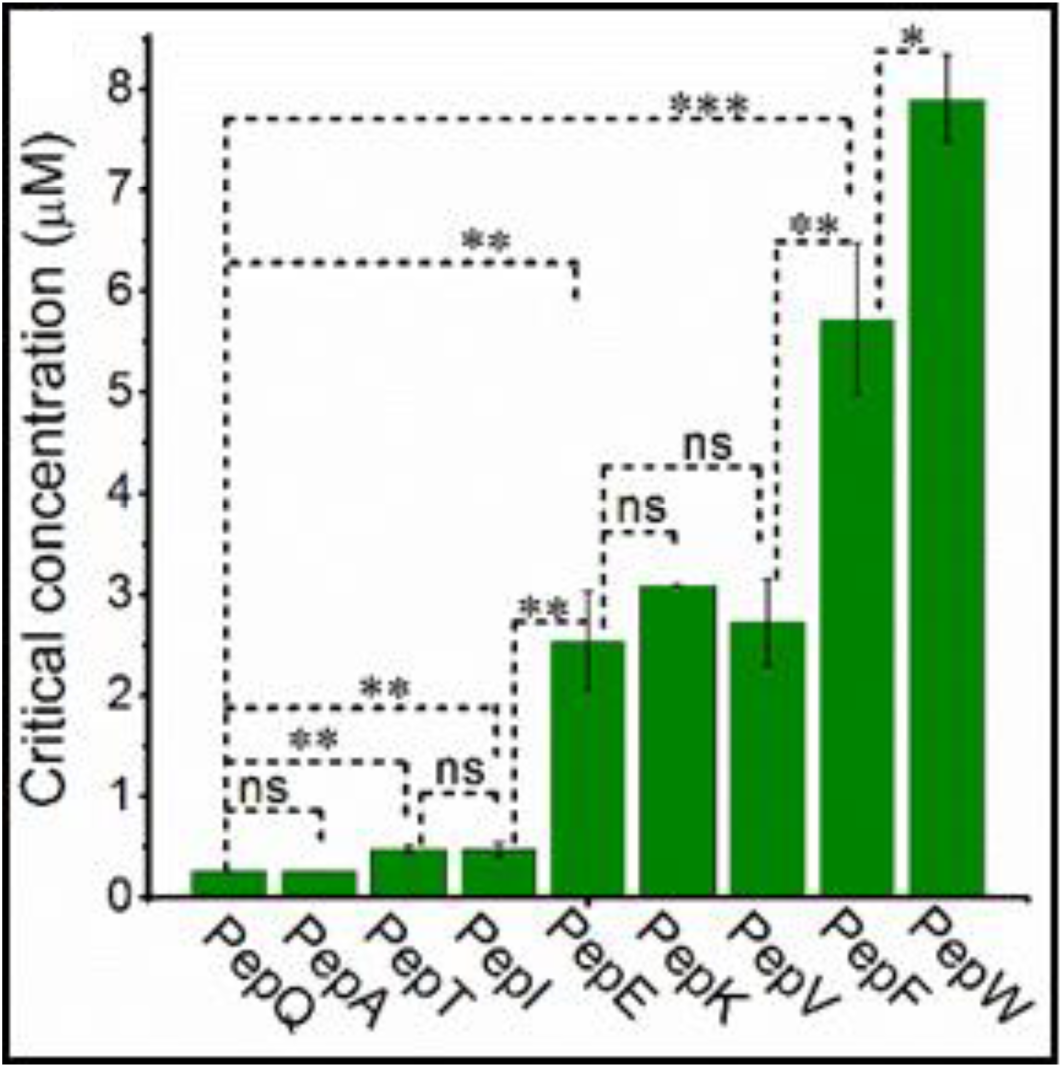
Critical concentrations (*Cr*) values of PolyGln mutant peptides obtained from spontaneous and seeding aggregation kinetics in the forward direction and spontaneous reverse kinetics of aggregates in the backward direction. Statistical significance of measurement is represented by asterisks (***, P<0.0001; **, 0.001 < P < 0.01; *, 0.01 < P < 0.05; ns=not significant, P > 0.05). For all the experiments error bars represent standard error of mean for minimum n = 4

### PolyGln aggregation propensity scale of amino acids

Apart from understanding the stability/instability of PolyGln mutant fibers, caused by amino acid substitutions, the ΔG values (Table 3) were also used to calculate the free energy difference (ΔΔG) of all the peptides with respect to host, PepQ peptide. This generated an aggregation propensity scale of all the amino acids substituted in the host peptide PepQ (Gln 18^th^ position). Further this propensity scale was compared with the theoretical intrinsic aggregation propensity scale of amino acids (30). This theoretical scale was established by taking into account the hydrophobicity, α-helical propensity, β-sheet propensity and the net charge of an individual amino acid. Other model systems which were also compared with the PolyGln aggregation propensities includes PolyAla AAHP and Aβ_(1-40)_ peptide [Table 2, see footnote (d, e & f)]. The free energy differences (ΔΔG) with respect to Gln for all the mutants were found to be in the range of -0.02 kcal/mol to -2.12 kcal/mol (Table 2). ΔΔG values of peptide mutants in the fast aggregating group (PepA, PepT and PepI) were found close to each other, suggesting their similar aggregation propensity in PolyGln aggregation motif. ΔΔG value of PepE and PepV peptides of the intermediate aggregating group is ∼1.4 kcal/mol less than PepQ, indicating their moderate aggregation propensity. Similarly, ΔΔG values of slow aggregating mutants, PepF and PepW showed a difference of ∼2 kcal/mol from PepQ peptide. This data shows that Phe and Trp have the lowest aggregation propensity in PolyGln AAHP. Exceptionally, PepK ΔΔG value of ∼1.5 kcal/mol (Table 2) is close to the intermediate aggregating group, suggesting its propensity similar to that of Val and Glu. This could be attributed to the fact that charge residues are known to have similar poor aggregation propensity (16, 31, 32). So PepK might be exhibiting ΔG value close to that of PepE. Thus in addition to the large size of Lys, its charge may also be involved in slowing PepK aggregation in comparison to PepE (21).

**Table 2.** Comparison of aggregation and β-sheet propensities of amino acids in PolyGln AAHP with different aggregation and β-sheet model systems.

| Amino acid | <sup>a</sup>(Chou & Fasman) (24)<br>Pβ | <sup>b</sup>(Kim & Berg) (26) | <sup>c</sup>(Minor & Kim) (25) | <sup>d</sup>(Fandrich et al.) (7) | <sup>e</sup>(Blondelle et al.) (41) | <sup>f</sup>Intrinsic aggregation propensity scale (23) | <sup>g</sup>PolyGln AAHP |
| --- | --- | --- | --- | --- | --- | --- | --- |
| **Q** | 1.23 | -0.40 | 0.23 | -0.51 | 0 | -6.00 | 0.00 |
| **A** | 0.97 | -0.35 | 0 | 0.41 | 75 | -3.31 | -0.02 |
| **I** | 1.60 | -0.56 | 1.0 | -0.68 | 0 | 0.93 | -0.38 |
| **T** | 1.20 | -0.48 | 1.10 | 0.11 | ~87 | -2.12 | -0.43 |
| **E** | 0.26 | -0.41 | 0.01 | 0.64 | ~20 | -10.38 | -1.42 |
| **V** | 1.65 | -0.53 | 0.82 | -0.03 | 50 | 0.49 | -1.47 |
| **K** | 0.74 | -0.41 | 0.27 | 1.27 | 0 | -9.55 | -1.54 |
| **F** | 1.28 | -0.56 | 0.86 | -0.21 | 50 | 2.80 | -1.93 |
| **W** | 1.19 | -0.48 | 0.54 | -0.17 | ~60 | 2.92 | -2.12 |
| **P** | 0.62 | -0.23 | <-3 | 1.02 | 0 | -11.96 | ND |

\*a: P $\beta$ (Conformation parameter of amino acids given by statistical analysis), $R^2 = 0.02$ ; b: $\Delta\Delta G$ (kcal/mol), given by using Zn finger peptide, $R^2 = 0.06$ ; c: $\Delta\Delta G$ (kcal/mol) given by using IgG binding domain of protein G, $R^2 = 0.01$ ; d: $\Delta\Delta G$ (kcal/mol) given by using A $\beta_{(1-40)}$ , $R^2 = 0.01$ ; e: % aggregation, given by using PolyAla peptide model, $R^2 = 0.002$ ; f: Intrinsic aggregation propensity $R^2 = 0.05$ ; g: $\Delta\Delta G$ (kcal/mol) from PolyGln peptide model; ND: not determined; $R^2$ : correlation coefficient of other models with respect to PolyGln AAHP. The data taken from the work of a, b, c, d & e authors were reproduced with due permission from the respective sources.

**Table 3.** Physicochemical parameters of amino acids incorporated in PolyGln mutants with their free energy (ΔG) of fibril formation.

| Amino acid | Hydrophobicity* | Size (VdWv) Å<sup>3</sup> | $\Delta G$ (kcal/mol) |
| --- | --- | --- | --- |
| **Q** | -3.5 | 114 | -9.36 ± 0.05 |
| **A** | 1.8 | 67 | -9.34 ± 0.05 |
| **I** | 4.5 | 124 | -8.98 ± 0.05 |
| **T** | -0.7 | 93 | -8.93 ± 0.32 |
| **E** | -3.5 | 109 | $-7.94 \pm 0.02$ |
| **V** | 4.2 | 105 | $-7.89 \pm 0.19$ |
| **K** | -3.9 | 135 | $-7.82 \pm 0.00$ |
| **F** | 2.8 | 135 | $-7.43 \pm 0.16$ |
| **W** | -0.9 | 163 | $-7.24 \pm 0.07$ |
| **P** | 1.6 | 90 | ND |

Physicochemical parameters are taken from reference (13) ;\*Negative values indicate nature of amino acid to be in hydrophilic environment and positive values denote their propensity to be in hydrophobic environment.
van der Waals volume (VdWv); ND ( not determined);
R<sup>2</sup>: correlation coefficient; R<sup>2</sup> between hydrophobicity and $\Delta G = 0.002$ ; R<sup>2</sup> between size of amino acid and $\Delta G = 0.8$ .

### PolyGln AAHP aggregation propensity scale versus scale from other amyloid models

Comparison of the ΔΔG values of amino acids obtained from PolyGln model system with the values calculated from other amyloid model systems, suggest a weak correlation among them [Table 2, see footnote (d, e & f)]. For example, R^2^ value of 0.05 was obtained by correlating PolyGln ΔΔG scale with theoretical aggregation propensity of amino acids (Figure 4A). Similarly, correlation of ΔΔG values of amino acids of PolyGln AAHP with the aggregation propensity scale of amino acids from Aβ_(1-40)_ peptide also showed poor R^2^ value of 0.01 (Figure 4B). Furthermore, comparison of ΔΔG values of PolyGln model system with aggregation tendency of amino acids in aggregation prone PolyAla AAHP repeat sequence also showed R^2^ = 0.002 (Figure 4C). Contrary to this, theoretical aggregation propensity of amino acids (except Gln) found to match significantly with those obtained from Aβ_(1-40)_ (12, 33) peptide showing R^2^ = 0.80 (Figure 4D).

**Figure 4.**
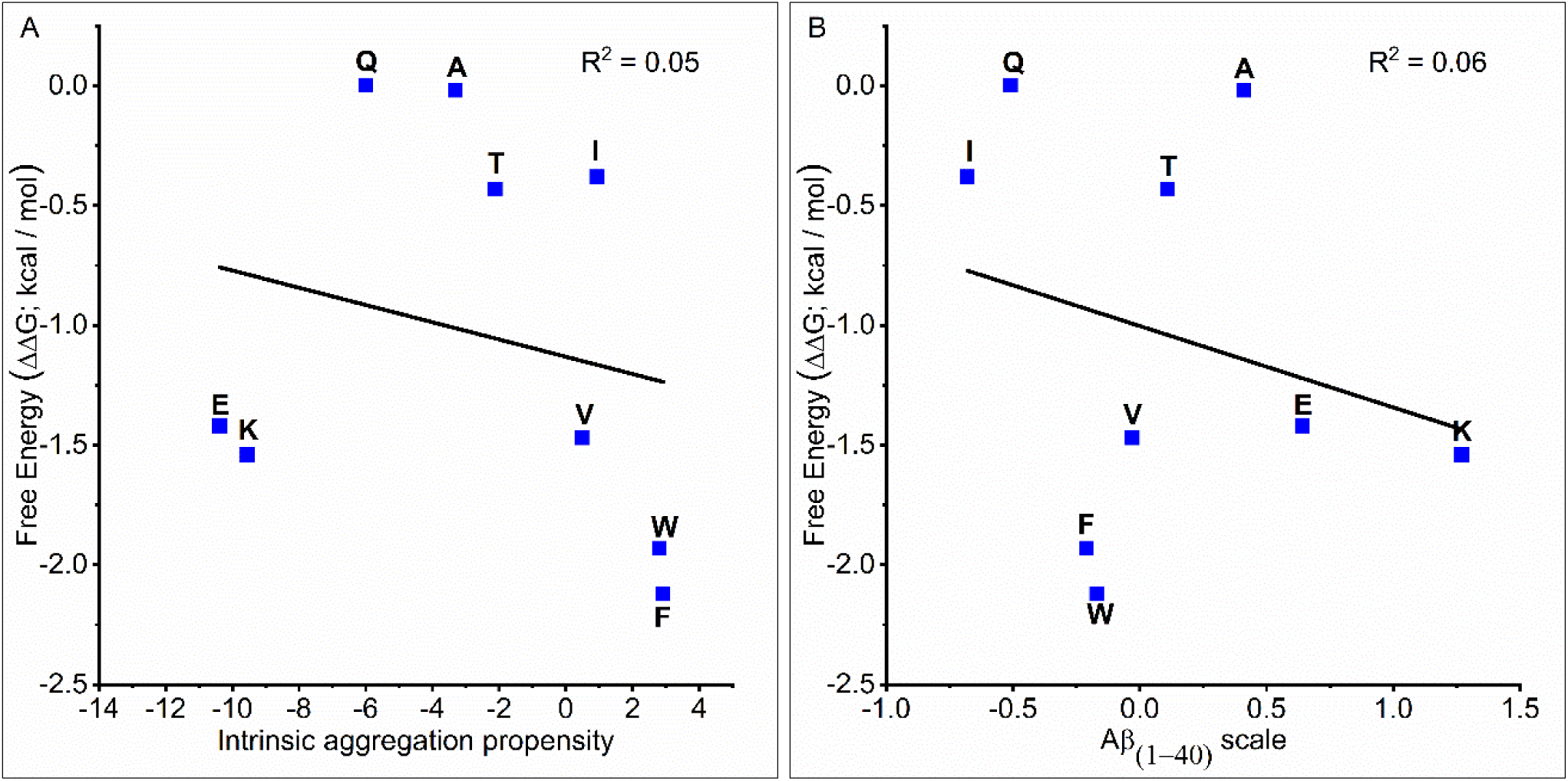
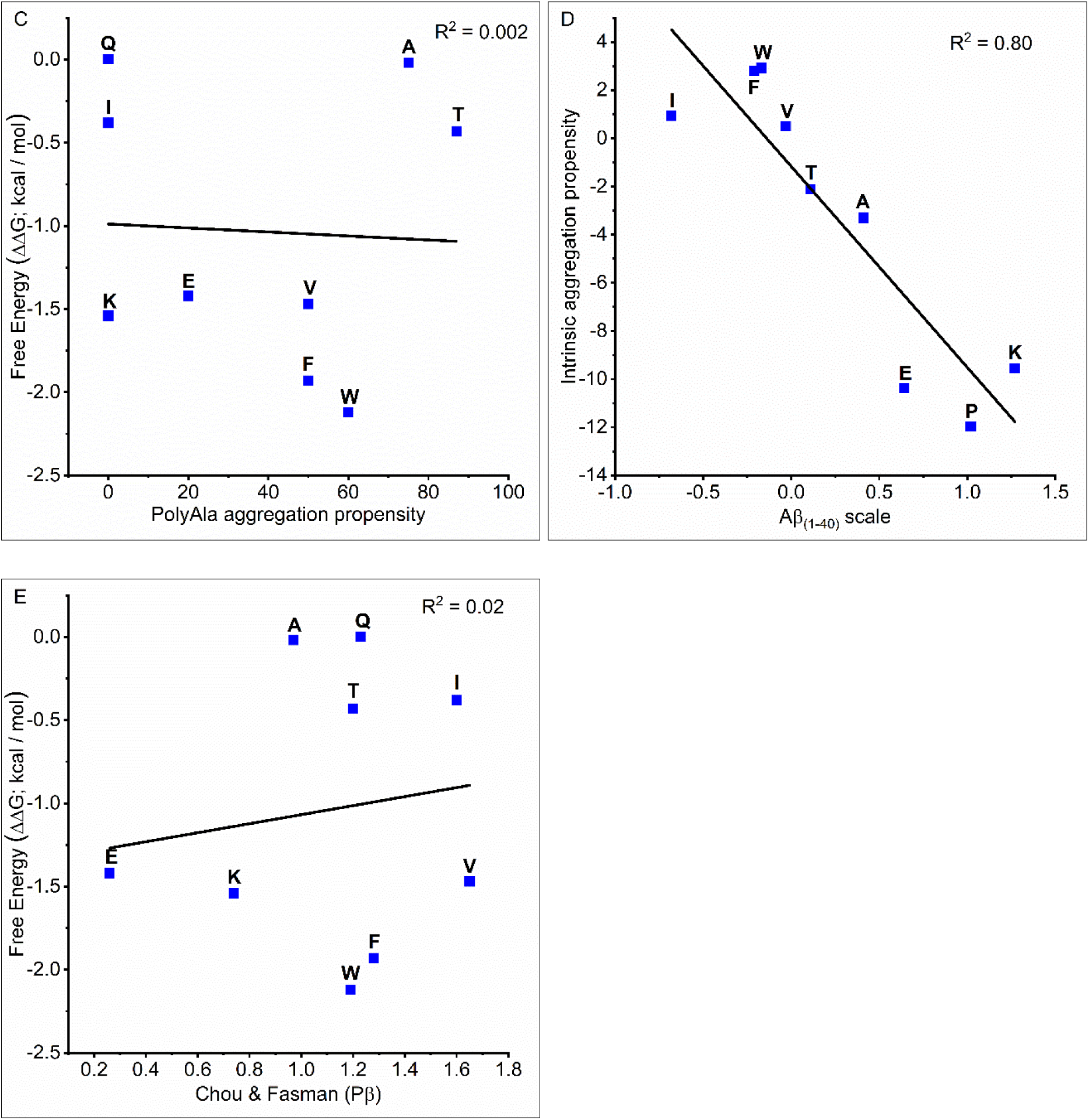
**A)** Correlation curve between ΔΔG of different amino acids from PolyGln AAHP and intrinsic aggregation propensity scale of amino acids (except Pro) R^2^ = 0.05, B) Correlation curve between ΔΔG of different amino acids from PolyGln AAHP and propensity scale of amino acid from Aβ _(1-40)_ peptide model (except Pro) R^2^ = 0.06, C) Correlation curve between ΔΔG of different amino acids from PolyGln AAHP and aggregation propensity scale of amino acids from PolyAla AAHP (except Pro) R^2^ = 0.002, D) Correlation curve between intrinsic aggregation propensity scale of amino acids and propensity scale of amino acids from Aβ _(1-40)_ peptide model except (Gln) R^2^ = 0.80 and E) Representative correlation curve between ΔΔG of different amino acids from PolyGln AAHP scale and Chou & Fasman β-sheet propensity scale, R^2^ = 0.02.

Different amino acids incorporated in PolyGln rich forty-six residue host peptide PepQ, showed impact on hierarchical levels of aggregation i.e. nucleation, overall aggregation rate, and fiber stability. Importantly, the aggregation propensity scale of amino acids in the PolyGln AAHP repeat system did not match with the theoretical intrinsic aggregation propensity, Aβ_(1-40)_ peptide and PolyAla AAHP sequence.

### PolyGln AAHP aggregation propensity scale versus β-sheet propensity scale from globular proteins

Since the PolyGln model system used in our study also promote the antiparallel β-sheet aggregating structure, it would be interesting to compare the ΔΔG values of amino acids from PolyGln model system with β-sheet propensity scale of amino acids that exists in β-sheet folding motifs of globular proteins. Strikingly, comparison of the ΔΔG values of amino acids obtained from the PolyGln model system with those calculated from other β-sheet model systems, also suggest a weak correlation among them (Table 2, see footnote a, b & c).

For example, the R^2^ value obtained by correlating conformational parameters of amino acids given by Chou & Fasman (16) was 0.02 (Figure 4E). Similarly, R^2^ of 0.06 was obtained by comparing ΔΔG values with Zn finger peptide (32). Comparison of ΔΔG values with IgG binding domain of protein G showed R^2^ = 0.01. This analysis confirms that the differential β-sheet propensity of amino acids exists in β-sheet folding motifs of globular proteins, Aβ peptide model and β-sheet rich aggregates of AAHP repeats. Contrary to this, β-sheet propensity of amino acids in globular proteins and Aβ_(1-40)_ (12, 33) peptide matches significantly with R^2^ = 0.86.

The above data suggests that hydrophobicity, net charge reduction of protein and intrinsic β-sheet propensity of amino acids that promote aggregation in globular proteins and amyloids may not govern the folding and aggregation of PolyGln AAHP repeats. To confirm it, hydrophobicity and size of various amino acids (van der Waals volume) substituted in PolyGln AAHP were fitted against the free energy of fibril formation (ΔG; Table 3).

### PolyGln ΔG scale comparison versus hydrophobicity and van der Waals volume of amino acids

Hydrophobicity did not correlate well with PolyGln free energy scale (ΔG); R^2^ = 0.001 (Figure S3), but good correlation (R^2^ = 0.8) was obtained when the van der Waals volume of amino acids was compared with the ΔG values (Figure 5C). It is important to note that correlation of 0.8 was obtained using seven amino acids only and does not contain Pro, Gln and Ile. When these three amino acids were included in the correlation of van der Waals volume and ΔG, R^2^ value became 0.1 (Figure 5C). On removing Pro and keeping both Gln and Ile along with seven amino acids, the R^2^ value becomes 0.1. Excluding Pro with either Gln or Ile the R^2^ value became ∼0.3 (Figure S4). Thus, Pro behaves as a major disruptor of the correlation. It suggests that van der Waals volume of Pro does not play a role in the non-aggregating nature of PepP. The presence of pyrrolidine ring structure in the Pro prevents its participation in the H-bonding between NH and CO groups of other amino acids (34, 35). The presence of the ring also confronts its presence in the β-sheet because the ring restricts its φ angle to -60 (34). This makes PepP non-aggregating in the amyloid core structure. Small deviation of Ile and Gln from the correlation could be due to their long side chain containing four carbon atoms. Gln being in PepQ, aggregation of it will not only be dictated by its van der Waals volume, but also through side chain hydrogen bond formation capability(20). Ile has β-branching due to which it may fit in the β-sheet folding motif of PolyGln in a similar fashion as that of Gln.

**Figure 5.**
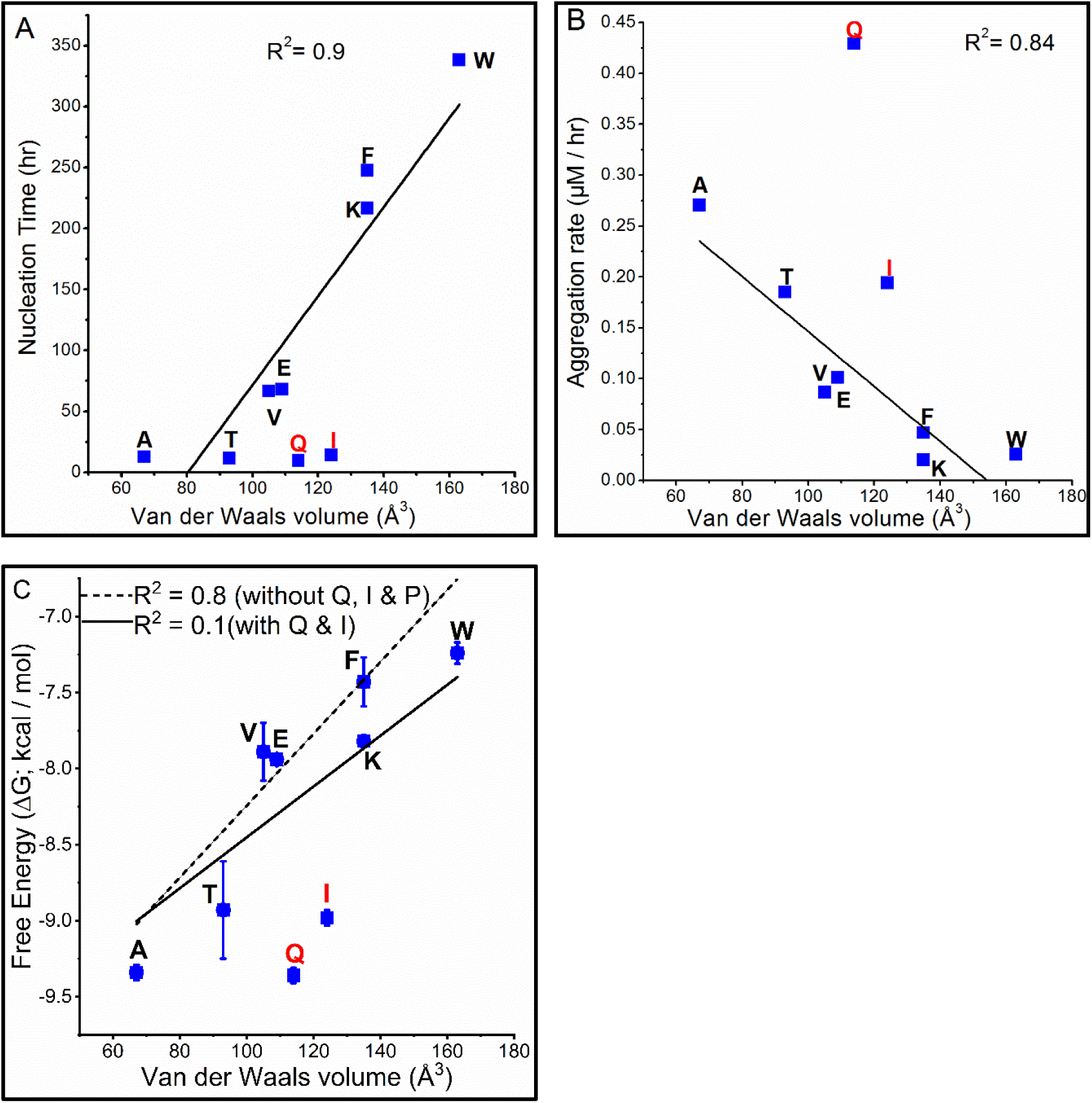
Correlation curves between van der Waals volume of amino acids and different biophysical parameters A) Nucleation time (hr), B) Aggregation rate (μM/hr) and C) Free energy (kcal/mol). R^2^ values are reported with the exception of Gln, Ile and Pro amino acids (see the discussion) while in figure 5C R^2^ values comparison is shown. Error bars represent standard error of mean.

## Discussion

Differential aggregation kinetics data obtained from insertion of various amino acids in the antiparallel β-sheet model peptide suggest that it is a reasonable model to carry out the mutational experiment. The data presented in this study delivers a major outcome i.e. aggregation and β-sheet propensity of the substituted amino acids using PolyGln AAHP model system are different from globular proteins, amyloid models and PolyAla AAHP system.

PolyGln peptides containing amino acids with high β-sheet propensity: Ile (PepI), Thr (PepT), Val (PepV), Phe (PepF) and Trp (PepW) were slow in aggregation as compared to Gln containing PepQ host peptide (Figure 2B). It means that amino acids having high β-sheet propensity in globular and other amyloid systems do not fit appropriately in the PolyGln β-sheet aggregation motif (Table 2). Moreover, PepA contains Ala with low β-sheet propensity in globular proteins, aggregate similar to PepQ (Figure 2B). Similarly, both Glu and Lys have very low β-sheet propensity but PepE aggregates faster than PepK.

Hydrophobic amino acids: Ile, Val, Phe and Trp are known to have high aggregation propensity in globular proteins (12, 16, 31, 32, 36). Mutational analysis from different intrinsically disordered amyloid forming systems, suggest that specific hydrophobic stretches in Aβ_1-40_ (10), α-synuclein (37), τ fragment from PHF43 alone (38), are capable of fibrillogenic aggregation. Deletion or substitution of these amino acids can suppress their aggregation capability towards amyloid formation. In support of it, acylphosphatase (AcP), a globular protein, exhibits similar results in amyloid formation, indicating the importance of hydrophobic residues (13, 14). Hydrophobic stretches can act as nucleation points to initiate the aggregation process and adjust well in the β-sheet rich structure (39). Interestingly, PolyGln peptides are intrinsically disordered, hydrophilic and do not contain any special hydrophobic stretch. As compared to other model systems, our results in PolyGln AAHP peptide model showed that Trp, Phe, Val and to some extent Ile, have less tendency to adopt in this β-sheet structure while aggregation. Another possibility for the slow aggregation behavior of the peptides containing hydrophobic residues is their presence in the hydrophilic microenvironment of Gln residues (Figure 2B). While these residues have high β-sheet propensity in native proteins and other amyloid systems, they may not be adjusting well due to exclusion from the PolyGln motif or causing minor structural changes around PolyGlns. Contrary to this Val and Ile are the two most hydrophobic amino acids (Table 1) and both PepV and PepI were aggregating much faster than PepW and PepF. In support of this, Ala which is also hydrophobic and has low β-sheet propensity, PepA is the second fast peptide mutant to aggregate. In a similar context Thr is hydrophilic and polar which is supposed to get easily adjusted in the PolyGln folding motif but PepT appeared to be the third fast aggregating mutant (Figure 2A). These points emphasize that hydrophilicity/hydrophobicity of amino acids is not playing a major role in controlling the aggregation behavior of these PolyGln mutant peptides.

Charge is another important determinant of aggregation behavior of amyloid forming proteins (14). High net charge on the proteins/peptides prevents self-association of their monomers to aggregation (15). Amino acid substitution in the AcP protein showed that higher net charge, decreases its aggregation propensity. Charge residues are present on the edge of β-strands of globular proteins to suppress aggregation (40). Similarly in GFP protein (41) and KTVIIE peptide (42), charge neutralization enhances their aggregation tendency. Lys and Glu charged amino acids have low β-sheet propensity but Lys is better compatible than Glu in a β-sheet model system (31, 32, 36). In our study charged residues delayed the nucleation time (Figure 2C), but Lys has extended it more as compared to Glu. One can possibly argue that in PepE where net charge on peptide is +3 would aggregate faster than PepK where net charge is +5. If that is the case, then PepE should have been even fast aggregating than PepQ which has +4 charge (Table 1, peptide design). Likewise, PepK due to +5 charge should be the slowest aggregating peptide. Rather PepF and PepW aggregates at somewhat similar rates as that of PepK (Figure 2B). This discussion along with the earlier discussion on hydrophobicity further confirm that hydrophobicity, charge and β-sheet propensity of amino acids, do not play a direct role in controlling the aggregation behavior of PolyGln AAHP peptides.

Such an unusual behavior of different amino acids in the PolyGln β-sheet motif is also supported by PolyGln amyloid core structure (20). According to this solid-state NMR structure, PolyGln aggregates contain antiparallel β-sheets having intramolecular β-hairpins. This structure suggests that the distance within two β-strands in a given antiparallel β-sheet is < 6.5 Å. Based on this interstrand distance, it can be predicted that amino acids having van der Waals volume < 143.7 Å^3^ (4/3 × *π* × (6.5/2)^3^) are more likely fit in the PolyGln β-sheet motif. As per this calculation, amino acids used in our study except Trp (163 Å^3^), van der Waals volume for all of them ranges within 67 Å^3^ to 135 Å^3^ and therefore they should fit well in the PolyGln β-sheet core. Our correlation results obtained between van der Waals volume and nucleation time, aggregation rate and free energy of fibril formation suggests a gradual size dependent trend. PepA and PepT with smaller amino acids Ala (67 Å^3^) and Thr (93 Å^3^) respectively, showed less nucleation time and high aggregation rate and fibril stability. Similarly, PepV and PepE with Val (105 Å^3^) and Glu (109 Å^3^) showed moderate nucleation time and aggregation rates and fibril stability. Furthermore, larger amino acids containing PepK (Lys; 135 Å^3^), PepF (Phe; 135 Å^3^) and PepW (Trp; 163 Å^3^) were slow aggregating with larger nucleation time and less thermodynamic stability. It is also possible that in comparison to our PolyGln model system, the interstrand distance limit observed for uninterrupted PolyGln < 6.5 Å is slightly less. It is because in our PolyGln peptide model Pro-Gly a turn inducing pair is present at regular intervals which prevents the sliding of Gln chains and helps in maintaining proper distance between two β-strands. Based on the presence of β-hairpin structure in the PolyGln amyloid core (20) and also during nucleation of PolyGln peptides (43, 44) it can be stated that the residues causing any sort of hindrance, can cause delay in the nucleation and overall aggregation rate of PolyGln AAHP. In this context the correlation observed between van der Waals volume of amino acids and nucleation time, aggregation rate and thermodynamic stability of fibril formation fits well.

To further strengthen our results that correlation results are indeed size dependent and other physicochemical parameters such as hydrophobicity and charge are not playing any major role in governing the PolyGln aggregation, we further examined these parameters against the aggregation behavior of PolyGln mutant peptides. However, Pro, Ile and Gln did not fit well in the correlation. Pro (90 Å^3^) being the β-sheet breaker distorted the correlation to greater extent. Throughout the proteome, Pro is known as a secondary structure breaker. Thus, despite having smaller size, PepP did not show any aggregation (Table 3). Therefore, the non-aggregating nature of PepP does not depend on its van der Waals volume. Ile amino acid (124 Å^3^ size) is larger than Glu (109 Å^3^) and Val (105 Å^3^), but PepI aggregates faster than both PepE and PepV. This could be due to the β-branched chain of Ile which can adjust itself easily in the PolyGln β-sheet motif as compared to the Glu and Val. Gln has a size of 114 Å^3^ but still aggregates exceptionally faster than its counterparts with small sizes like Thr (93 Å^3^), Glu (109 Å^3^) and Val (105 Å^3^) (Figure 2B). It could be due to the extremely efficient H-bonding capability of Gln (45) within its same background as a host (20). ssNMR study of aggregates from Huntington exon 1 fragment and different PolyGln peptides showed a network of common interdigitated extended and rigid Gln side chains in the PolyGln amyloid core (20). These results also suggest that Gln is capable of making backbone– backbone, backbone–side chain and side chain–side chain hydrogen bond interactions and this tight H-bonding among Glns is very difficult to replace by any other amino acid (46). This overall mutational analysis highlight that van der Waals volume (VdWv) of amino acids showed positive correlation with nucleation time and thermodynamic stability of different peptides and inverse correlation with the aggregation rate (Figure 5).

## Conclusion

Similarity between aggregation and β-sheet propensity of amino acids among amyloids and native proteins is well documented (12) (Table 2). This suggests a cooperative behavior of neighboring residues adjusting themselves according to the amino acid present in between them or vice-versa (12). In this study we showed that these rules are not the same in PolyGln AAHP. In the similar sequence background, the rules of aggregation and β-sheet propensity of amino acids differ not only from globular proteins and other amyloid systems but also among other AAHP repeats like PolyAla. Notably, the differential aggregation behavior of PolyGln peptides only correlated with van der Waals volume of the incorporated amino acid and not with any other physicochemical characteristic. Elucidating the rules that govern an AAHP repeat sequence to fold, unfold and form amyloid-like structure may help in decoding their conformation, stability and structure-function relationships under cellular conditions. AAHP repeats of all the amino acids are present in the human proteome (28, 56). Interestingly, depending on the association with various diseases only three AAHP repeats i.e. PolyGln (27), PolyAla (57) and more recently PolyAsn (58) have been studied till date. The same host-guest approach could be taken as a starting point towards establishing the principles governing their folding or aggregation. Human sequence database of PolyGln proteins underscore the presence of several interruptions in them (28) (Table S3). However, the exact relevance of these interruptions except few (31, 59) in the disease context is still not known. The outcome of this study can be utilized and replicated in such PolyGln protein sequences to understand the impact of any interruptions. Interestingly, amino acid interruptions in PolyGln sequences are not only capable of modulating their aggregation behavior (32, 60) but also its aggregate structure (60) and toxicity (54, 59). Homorepeats and pattern database (HRaP) (56) revealed the presence of different amino acid repeats with variable length patterns across 122 different proteomes. Based on such a wide occurrence of these repeat sequences, the present study will open new avenues of research in testing the structural role of different stretches of PolyGln and other AAHP sequences in proteins of prokaryotes and eukaryotes. This study also paves the way to tune the fiber formation pathways on the basis of amino acid inserted in AAHP of PolyGln and other amino acid stretches to form multitude of amyloid structures, having implications in biology and design of nanomaterials.

## Acknowledgements

AKT acknowledges Department of Biotechnology, Government of India (Grant No: BT/PR3041/NNT/545/2011) for financial support. RM thanks IIT Kanpur for providing Ph. D fellowship.

## Supplementary Information

### MATERIALS AND METHODS

#### (M1) Peptide synthesis and solubilization

The peptides were synthesized by solid phase method and obtained in crude form, from Keck Biotechnology Centre at Yale University. These were purified using reverse-phase HPLC, lyophilized and stored at -80 °C. Purified peptides were then disaggregated in 1:1 ratio of Trifluoroacetic acid (TFA) and *Hexafluoroisopropanol (HFIP) (Sigma-Aldrich).TFA and HFIP solvents were evaporated using gentle stream of nitrogen gas. The glass vials were desiccated for two hours to remove any remaining volatile solvents, dissolved using water-TFA pH 3 and centrifuged* (Thermo Scientific Sorvall MTX 150 Micro-ultracentrifuge) in polypropylene micro vials, using S80AT2 rotor at 25000 g *for four hours. This ensured pelleting of undissolved peptide material. Supernatant containing soluble peptide fraction was collected gently for concentration determination and setting up of the aggregation reaction (1).*

#### *(M2)* Spontaneous aggregation kinetics by RP-HPLC sedimentation assay

Spontaneous aggregation reaction of all PolyGln peptides were monitored by RP-HPLC (Agilent 1260 infinity instrument, propylene-bridged bidentate-C18 silane column, particle size 3.5 μM, pressure limit 600 bar, flow rate 1 ml/min). For setting up the spontaneous aggregation reaction, desired amount of peptide after disaggregation and ultracentrifugation (section M1) was mixed with 10X PBS (pH 7.2) and final reaction volume was made up using water-TFA pH 3 to achieve 1X PBS concentration. Sodium azide (0.05 %) was added to the reaction mixture to prevent microbial contamination. Reaction mixture was kept at 37 °C for aggregation in an incubator (Mahendra Scientific Instruments)(1). To monitor the aggregation kinetics, an aliquot of an on-going reaction was ultracentrifuged for 30 minutes at 25000 g and supernatant concentration was measured at 215 nm by RP-HPLC using standard curve obtained for each peptide (1). All the peptides show differential hydrophobicity, thus yielded different retention times. Aggregation rates (μM/hr) were obtained by calculating the slope of the curve fitted between monomer concentration and time as shown in the Figure S1.

#### (M3) Preparation of standard curves to determine concentration of PolyGln peptides by RP-HPLC method

To generate a standard curve, the concentration of the disaggregated peptide (section M1) stock solution was measured using UV spectrophotometer (Perkin Elmer) at 215 nm which corresponds to the absorbance of peptide bond. The absolute concentration of peptide stock was obtained using Beer-Lambert law: 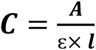 where [C = molar concentration of peptide, A = absorbance, ε = molar extinction coefficient of peptide and l = path length (1 cm)].

Molar extinction coefficient was calculated for each peptide mutant using the formula given in the paper (Kuipers and Gruppen, 2007) (2) (Table M1). The following equation was used:

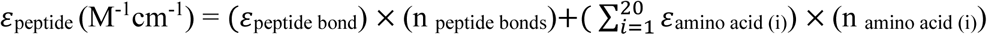

where ε_peptide_ is the molar extinction coefficient of peptide, ε_peptide bond_ is the molar extinction coefficient of peptide bond, n_peptide bonds_ is the number of peptide bonds in the peptide, ε_amino acid_ is the molar extinction coefficient of amino acid and n_amino acid_ is the number of amino acids in the peptide. Molar extinction coefficient values suggest that all the peptides have almost similar values with the exception of PepF and PepW.

Thereafter, the peptide stock solution is serially diluted to 2, 4, 8, 16 and 32 folds. The absorbance of these diluted peptide fractions were determined using UV spectrophotometer (Perkin Elmer) at 215 nm and their concentration was measured using Beer-Lambert law.

**Table M1.** Molar extinction coefficients and standard curve equations of different peptides.

| Peptide | Molar extinction coefficient ($\epsilon$)(M-1 cm-1) | Standard curve equation, 'y' is peak area and 'x' is $\mu\text{g}$ of peptide |
| --- | --- | --- |
| PepQ | 54836 | $y = 9 \times 10^6 x - 1 \times 10^6$ |
| PepA | 54726 | $y = 9 \times 10^6 x - 1 \times 10^6$ |
| PepT | 54735 | $y = 9 \times 10^6 x - 1 \times 10^6$ |
| PepI | 54739 | $y = 9 \times 10^6 x - 1 \times 10^6$ |
| PepE | 54772 | $y = 9 \times 10^6 x - 1 \times 10^6$ |
| PepV | 54737 | $y = 9 \times 10^6 x - 1 \times 10^6$ |
| PepK | 54735 | $y = 9 \times 10^6 x - 1 \times 10^6$ |
| PepF | 59894 | $y = 9 \times 10^6 x - 2 \times 10^6$ |
| PepW | 83744 | $y = 1 \times 10^7 x - 574545$ |
| PepP | 57369 | $y = 9 \times 10^6 x - 1 \times 10^6$ |

After getting the concentration values, different volumes (μl) of these diluted peptide stocks were injected into HPLC and their corresponding peak areas from their respective chromatograms were calculated. A peak area in the chromatogram is directly proportional to the amount of peptide injected into the HPLC column. Thus standard curve was plotted between peak areas (Y axis) *vs.* μg (X axis) of the peptide injected and the values were subjected to linear curve fitting. The linear fit provided a straight line equation (Table M1) which was used for calculating the unknown peptide concentration.

#### (M4) Measurement of critical concentration (*Cr*) of PolyGln peptide mutants

*Cr* values of all the PolyGln mutants were calculated by spontaneous and seeding aggregation kinetics. RP-HPLC with ∼200 ng limit of detection was used. *Cr* values of peptides were also measured reversibly to establish the presence of dynamic equilibrium (3). A fraction of sample at the end of aggregation reaction was diluted in fresh 1X PBS pH 7.2 buffer such that the final concentration of peptide (aggregates) in the solution is above its corresponding *Cr* obtained from the forward reaction. After dilution it was kept at 37 °C and release of monomer from the aggregates was monitored using RP-HPLC method as described in method M2. Conversion of *Cr* to thermodynamic parameter 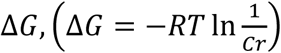, allowed the measurement and comparison of the thermodynamic stability of aggregates of different PolyGln mutants (4, 5).

#### (M5) Comparison of aggregation propensity of amino acids in PolyGln amino-acid-homopolymer repeat system and other models

Free energy value (ΔG) of fibril formation from different PolyGln mutants were calculated as described in method M4. The free energy values difference (ΔΔG) was calculated to compare it with aggregation and β-sheet propensities from other model systems as represented in Table 2. ΔΔG (kcal/mole) is the difference in free energy of fibril formation (ΔG) for each substituted amino acid residue relative to Gln at 18^th^ position (Table 1). For example (ΔG for PepQ = -9.36 kcal/mol) – (ΔG for PepA = -9.34 kcal/mol) which gives the expression ΔΔG_(Ala)_ = -0.02 kcal/mol (Table 2). Similar method is used for calculating ΔΔG values of all other peptides. Previously, β-sheet conformational parameter given by Chou & Fasman was compared with the ΔΔG values of substituted amino acids from globular proteins and other amyloid systems showing good correlation among them (6-8). Similar comparison of the ΔΔG values obtained for amino acids from PolyGln peptide model was carried out with the β-sheet propensity scale from Chou & Fasman (6), Kim & Berg (7), Minor & Kim (8), Fandrich *et al.*(9) and Blondelle *et al.*(10) model systems (Table 2). The correlation coefficient obtained between different scales and PolyGln scale is shown in the footnote of Table 2.

**Figure S1.**
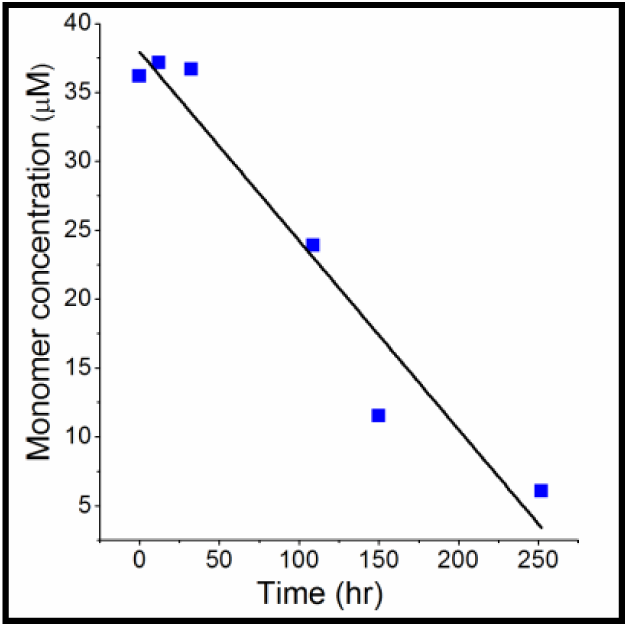
Representative spontaneous aggregation kinetics curve, showing aggregation rate of a peptide. Aggregation data is subjected to linear curve fitting between concentration (μM) and time (hr).The straight line obtained is *y* = −0.137*x* + 37.934, where slope *x* is the aggregation rate (μM/hr).

**Figure S2.**
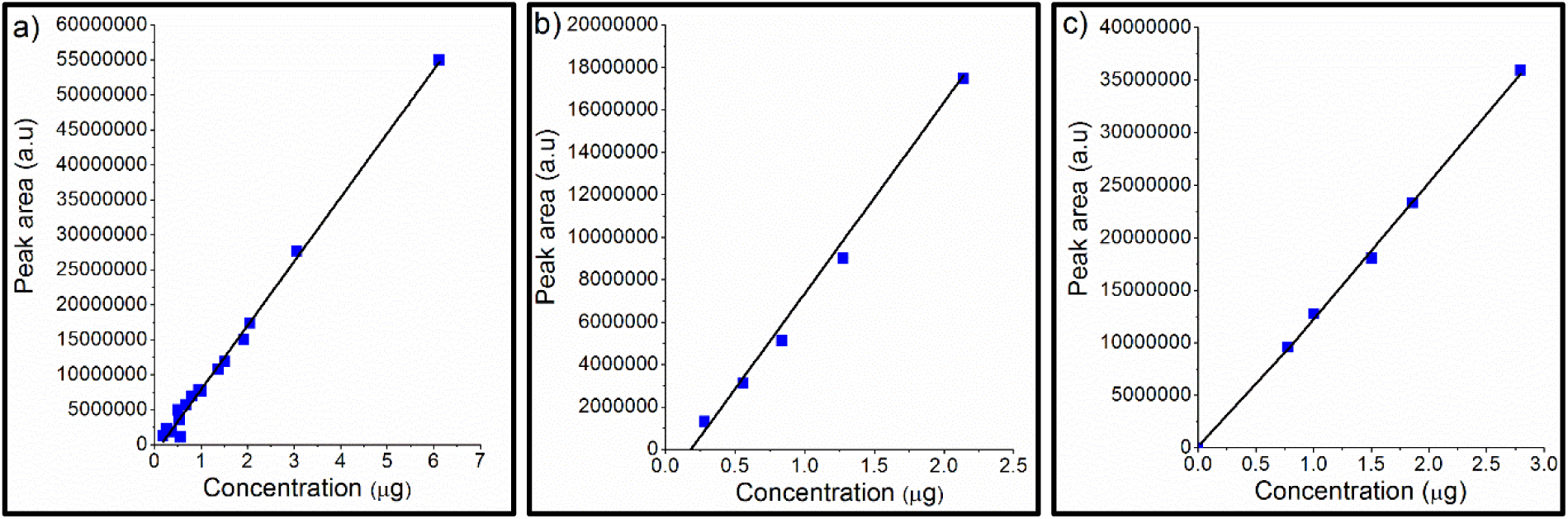
Representative standard curves of peptides between different amounts *vs* corresponding areas obtained in RP-HPLC at 215 nm. a) PepQ b) PepF and c) PepW.

**Table S1.**
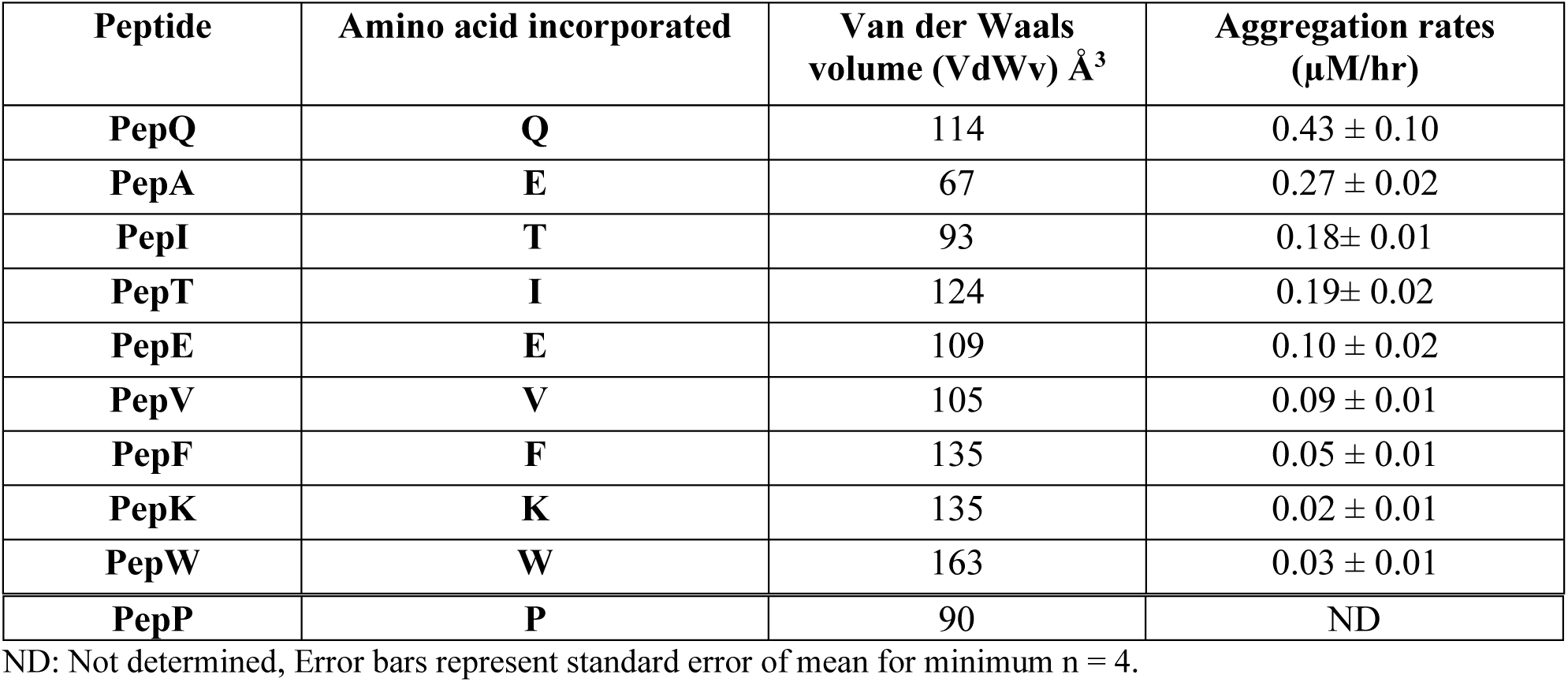
Comparison of spontaneous aggregation rates of different PolyGln peptides.

| Peptide | Amino acid incorporated | Van der Waals volume (VdWv) $\text{\AA}^3$ | Aggregation rates ($\mu\text{M/hr}$) |
| --- | --- | --- | --- |
| PepQ | Q | 114 | $0.43 \pm 0.10$ |
| PepA | E | 67 | $0.27 \pm 0.02$ |
| PepI | T | 93 | $0.18 \pm 0.01$ |
| PepT | I | 124 | $0.19 \pm 0.02$ |
| PepE | E | 109 | $0.10 \pm 0.02$ |
| PepV | V | 105 | $0.09 \pm 0.01$ |
| PepF | F | 135 | $0.05 \pm 0.01$ |
| PepK | K | 135 | $0.02 \pm 0.01$ |
| PepW | W | 163 | $0.03 \pm 0.01$ |
| **PepP** | **P** | 90 | ND |

ND: Not determined, Error bars represent standard error of mean for minimum n = 4.

**Table S2.** Critical concentrations of PolyGln peptides.

| Peptide | Amino acid incorporated | Van der Waals volume (VdWv) Å<sup>3</sup> | Cr (μM) |
| --- | --- | --- | --- |
| **PepQ** | **Q** | 114 | 0.25 ± 0.01 |
| **PepA** | **E** | 67 | 0.26 ± 0.01 |
| **PepI** | **T** | 93 | 0.47 ± 0.02 |
| **PepT** | **I** | 124 | 0.51 ± 0.13 |
| **PepE** | **E** | 109 | 2.54 ± 0.04 |
| **PepV** | **V** | 105 | 2.73 ± 0.42 |
| **PepF** | **F** | 135 | 5.73 ± 0.75 |
| **PepK** | **K** | 135 | 3.09 ± 0.01 |
| **PepW** | **W** | 163 | 7.90 ± 0.42 |
| **PepP** | **P** | 90 | ND |

\*ND: Not determined, Error bars represent standard error of mean for minimum n = 4.

**Figure S3.**
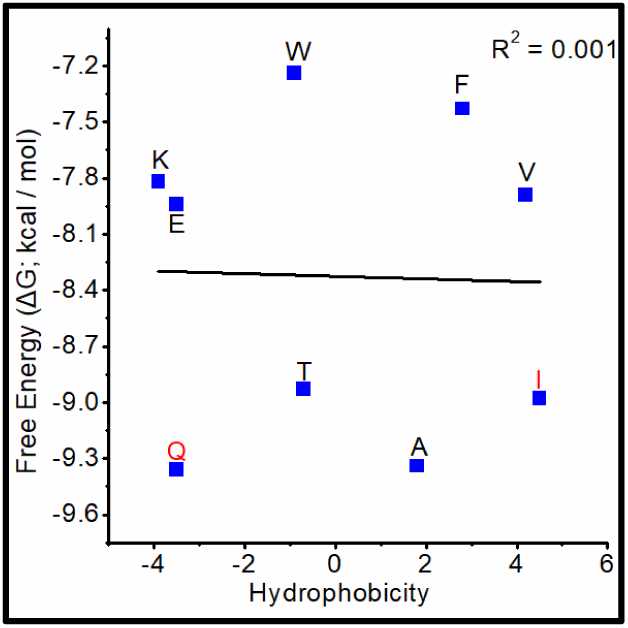
Correlation curve between ΔG of different amino acids [obtained from PolyGln AAHP scale] and hydrophobicity of amino acids.

**Figure S4.**
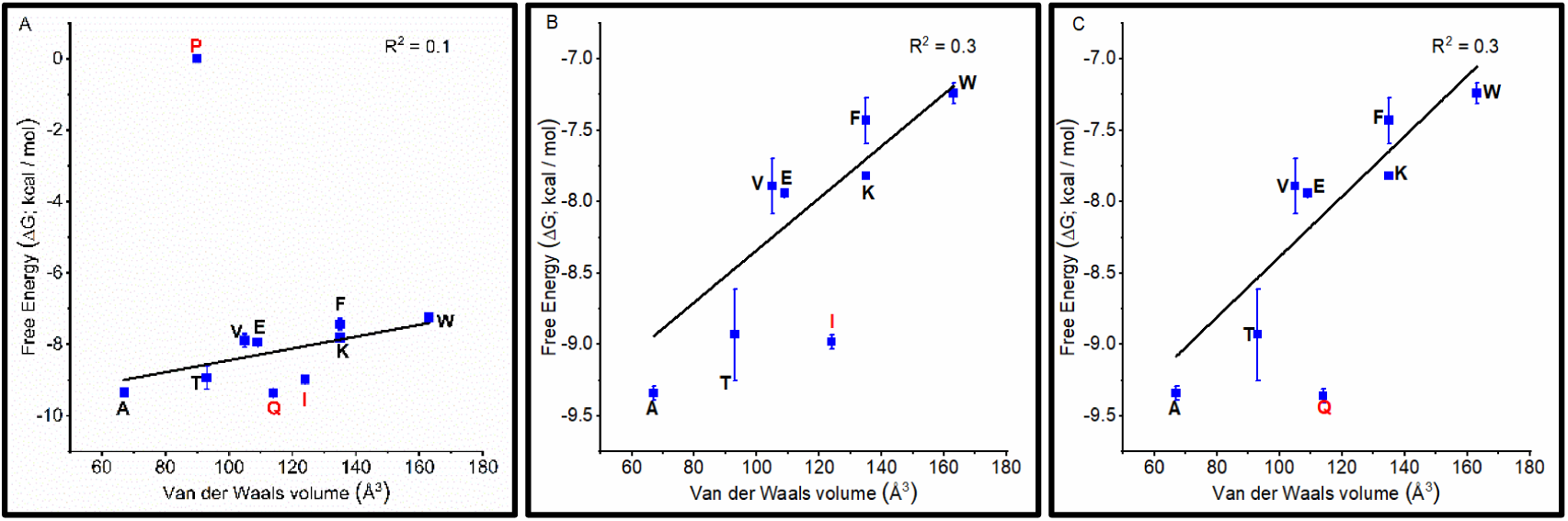
Correlation curve between ΔG of different amino acids [obtained from PolyGln AAHP scale] and their van der Waals volume: A) including Gln, Ile & Pro; R^2^ = 0.1, B) including only Ile; R^2^ = 0.3 and C) including only Gln; R^2^ = 0.3.

**Table S3.**
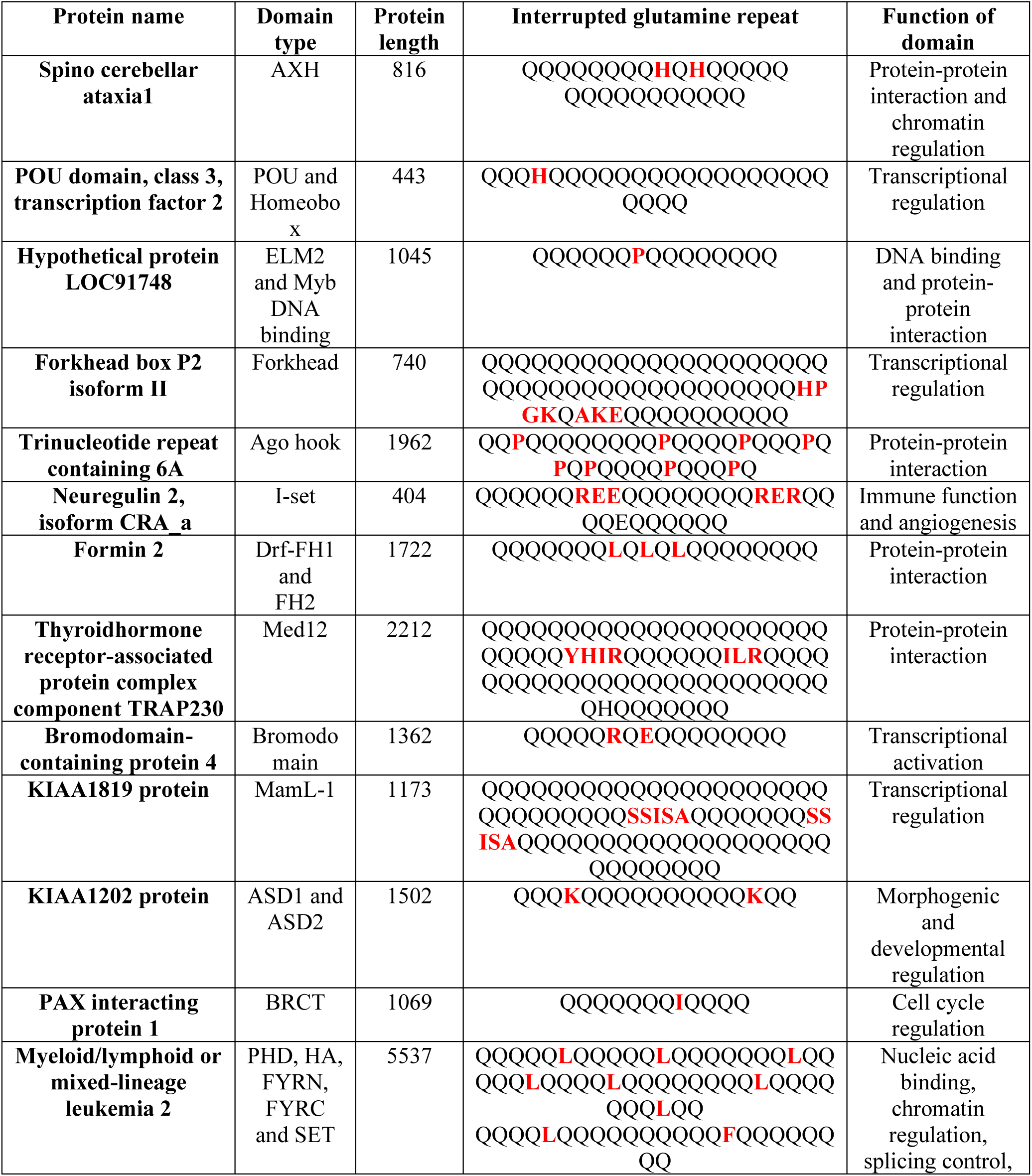
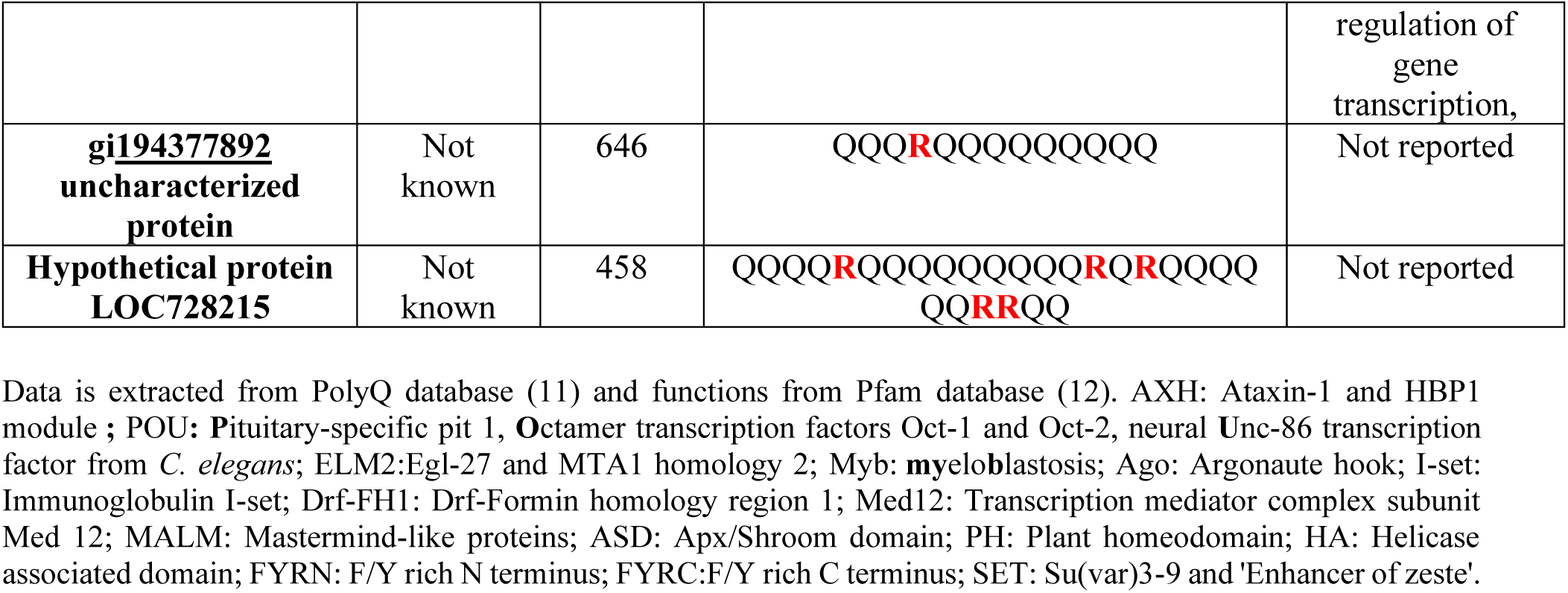
Some human proteins with interrupted PolyGln repeats.

## References

1. Lobanov MY, Sokolovskiy IV, & Galzitskaya OV (2013) HRaP: database of occurrence of HomoRepeats and patterns in proteomes. Nucleic acids research 42(D1):D273–D278.

2. Wetzel R (2012) Physical chemistry of polyglutamine: intriguing tales of a monotonous sequence. Journal of molecular biology 421(4-5):466–490.

3. Scherzinger E, et al. (1997) Huntingtin-encoded polyglutamine expansions form amyloid-like protein aggregates in vitro and in vivo. Cell 90(3):549–558.

4. Chen S, Berthelier V, Hamilton JB, O’Nuallai B, & Wetzel R (2002) Amyloid-like features of polyglutamine aggregates and their assembly kinetics. Biochemistry 41(23):7391–7399.

5. Chen S, Ferrone FA, & Wetzel R (2002) Huntington’s disease age-of-onset linked to polyglutamine aggregation nucleation. Proceedings of the National Academy of sciences 99(18):11884–11889.

6. Scherzinger E, et al. (1999) Self-assembly of polyglutamine-containing huntingtin fragments into amyloid-like fibrils: implications for Huntington’s disease pathology. Proceedings of the National Academy of Sciences 96(8):4604–4609.

7. Harper JD & Lansbury Jr PT (1997) Models of amyloid seeding in Alzheimer’s disease and scrapie: mechanistic truths and physiological consequences of the time-dependent solubility of amyloid proteins. Annual review of biochemistry 66(1):385–407.

8. Chiti F, et al. (1999) Designing conditions for in vitro formation of amyloid protofilaments and fibrils. Proceedings of the National Academy of Sciences 96(7):3590–3594.

9. Chiti F & Dobson CM (2006) Protein misfolding, functional amyloid, and human disease. Annu. Rev. Biochem. 75:333–366.

10. Williams AD, et al. (2004) Mapping Aβ amyloid fibril secondary structure using scanning proline mutagenesis. Journal of molecular biology 335(3):833–842.

11. Williams AD, Shivaprasad S, & Wetzel R (2006) Alanine Scanning Mutagenesis of Aβ(1-40) Amyloid Fibril Stability. Journal of Molecular Biology 357(4):1283–1294.

12. Hortschansky P, Christopeit T, Schroeckh V, & Fändrich M (2005) Thermodynamic analysis of the aggregation propensity of oxidized Alzheimer’s β-amyloid variants. Protein Sci 14(11):2915–2918.

13. Chiti F, et al. (1999) Mutational analysis of acylphosphatase suggests the importance of topology and contact order in protein folding. Nat Struct Mol Biol 6(11):1005–1009.

14. Chiti F, et al. (2000) Mutational analysis of the propensity for amyloid formation by a globular protein. The EMBO journal 19(7):1441–1449.

15. Chiti F, Stefani M, Taddei N, Ramponi G, & Dobson CM (2003) Rationalization of the effects of mutations on peptide andprotein aggregation rates. Nature 424(6950):805–808.

16. Chou PY & Fasman GD (1974) Conformational parameters for amino acids in helical, β-sheet, and random coil regions calculated from proteins. Biochemistry 13(2):211–222.

17. Street AG & Mayo SL (1999) Intrinsic β-sheet propensities result from van der Waals interactions between side chains and the local backbone. Proceedings of the National Academy of Sciences 96(16):9074–9076.

18. Thakur AK, et al. (2009) Polyglutamine disruption of the huntingtin exon 1 N terminus triggers a complex aggregation mechanism. Nature structural & molecular biology 16(4):380.

19. Thakur AK & Wetzel R (2002) Mutational analysis of the structural organization of polyglutamine aggregates. Proceedings of the National Academy of Sciences 99(26):17014–17019.

20. Hoop CL, et al. (2016) Huntingtin exon 1 fibrils feature an interdigitated β-hairpin–based polyglutamine core. Proceedings of the National Academy of Sciences 113(6):1546–1551.

21. Mishra R & Thakur AK (2015) Amyloid nanospheres from polyglutamine rich peptides: assemblage through an intermolecular salt bridge interaction. Org Biomol Chem 13(14):4155–4159.

22. O’Nuallain B, et al. (2006) Kinetics and Thermodynamics of Amyloid Assembly Using a High-Performance Liquid Chromatography–Based Sedimentation Assay. Methods in Enzymology, eds Indu K & Ronald W (Academic Press), Vol Volume 413, pp 34–74.

23. Jayaraman M KR, Wetzel R (2009) The impact of ataxin-1-like histidine insertions on polyglutamine aggregation. Protein Eng Des Sel. 22.(8):469–478.

24. Fernandez-Escamilla AM, Rousseau F, Schymkowitz J, & Serrano L (2004) Prediction of sequence-dependent and mutational effects on the aggregation of peptides and proteins. Nat Biotechnol 22(10):1302–1306.

25. Nakano M, Watanabe H, Starikov EB, Rothstein SM, & Tanaka S (2009) Mutation effects on structural stability of polyglutamine peptides by molecular dynamics simulation. Interdisciplinary sciences, computational life sciences 1(1):21–29.

26. Brummitt RK, Andrews JM, Jordan JL, Fernandez EJ, & Roberts CJ (2012) Thermodynamics of amyloid dissociation provide insights into aggregate stability regimes. Biophys Chem 168:10–18.

27. Cobb NJ, Apetri AC, & Surewicz WK (2008) Prion Protein Amyloid Formation under Native-like Conditions Involves Refolding of the C-terminal alpha-Helical Domain. J Biol Chem 283(50):34704–34711.

28. O’Nuallain B, Shivaprasad S, Kheterpal I, & Wetzel R (2005) Thermodynamics of Aβ(1-40) Amyloid Fibril Elongation†. Biochemistry 44(38):12709–12718.

29. Chen S, Berthelier V, Yang W, & Wetzel R (2001) Polyglutamine aggregation behavior in vitro supports a recruitment mechanism of cytotoxicity. Journal of molecular biology 311(1):173–182.

30. Pawar AP, et al. (2005) Prediction of “aggregation-prone” and “aggregation-susceptible” regions in proteins associated with neurodegenerative diseases. Journal of molecular biology 350(2):379–392.

31. Minor DL & Kim PS (1994) Measurement of the [beta]-sheet-forming propensities of amino acids. Nature 367(6464):660–663.

32. Kim CA & Berg JM (1993) Thermodynamic [beta] -sheet propensities measured using a zinc-finger host peptide. Nature 362(6417):267–270.

33. Christopeit T, et al. (2005) Mutagenic analysis of the nucleation propensity of oxidized Alzheimer’s beta-amyloid peptide. Protein Sci 14(8):2125–2131.

34. Creighton TE (1983) Proteins: Structure and Molecular Properties (Freeman, New York).

35. Wouters MA & Curmi PMG (1995) An analysis of side chain interactions and pair correlations within antiparallel β-sheets: The differences between backbone hydrogen-bonded and non-hydrogen-bonded residue pairs. Proteins: Structure, Function, and Bioinformatics 22(2):119–131.

36. Muñoz V & Serrano L (1994) Intrinsic secondary structure propensities of the amino acids, using statistical ϕ–ψ matrices: Comparison with experimental scales. Proteins: Structure, Function, and Bioinformatics 20(4):301–311.

37. Giasson BI, Murray IVJ, Trojanowski JQ, & Lee VMY (2001) A hydrophobic stretch of 12 amino acid residues in the middle of alpha-synuclein is essential for filament assembly. J Biol Chem 276(4):2380–2386.

38. Spillantini MG & Goedert M (2001) Tau gene mutations and tau pathology in frontotemporal dementia and parkinsonism linked to chromosome 17. Adv Exp Med Biol 487:21–37.

39. Chiti F (2006) Relative importance of hydrophobicity, net charge, and secondary structure propensities in protein aggregation. Protein Misfolding, Aggregation, and Conformational Diseases, (Springer), pp 43–59.

40. Richardson JS & Richardson DC (2002) Natural β-sheet proteins use negative design to avoid edge-to-edge aggregation. Proceedings of the National Academy of Sciences 99(5):2754–2759.

41. Raghunathan G, et al. (2013) Modulation of protein stability and aggregation properties by surface charge engineering. Mol Biosyst 9(9):2379–2389.

42. López de la Paz M, et al. (2002) De novo designed peptide-based amyloid fibrils. Proceedings of the National Academy of Sciences 99(25):16052–16057.

43. Kar K, et al. (2013) β-Hairpin-Mediated Nucleation of Polyglutamine Amyloid Formation. Journal of Molecular Biology 425(7):1183–1197.

44. Kar K, Jayaraman M, Sahoo B, Kodali R, & Wetzel R (2011) Critical nucleus size for disease-related polyglutamine aggregation is repeat-length dependent. Nature structural & molecular biology 18(3):328–336.

45. Rhys NH & Dougan L (2013) The emerging role of hydrogen bond interactions in polyglutamine structure, stability and association. Soft Matter 9(8):2359–2364.

46. Rhys N, Soper A, & Dougan L (2012) The hydrogen-bonding ability of the amino acid glutamine revealed by neutron diffraction experiments. The Journal of Physical Chemistry B 116(45):13308–13319.

## References

1. Chou PY & Fasman GD (1974) Conformational parameters for amino acids in helical, β-sheet, and random coil regions calculated from proteins. Biochemistry 13(2):211–222.

2. Kim CA & Berg JM (1993) Thermodynamic [beta] -sheet propensities measured using a zinc-finger host peptide. Nature 362(6417):267–270.

3. Minor DL & Kim PS (1994) Measurement of the [beta]-sheet-forming propensities of amino acids. Nature 367(6464):660–663.

4. Hortschansky P, Christopeit T, Schroeckh V, & Fändrich M (2005) Thermodynamic analysis of the aggregation propensity of oxidized Alzheimer’s β-amyloid variants. Protein Sci 14(11):2915–2918.

5. Blondelle SE, Forood B, Houghten RA, & Pérez-Payá E (1997) Polyalanine-based peptides as models for self-associated β-pleated-sheet complexes. Biochemistry 36(27):8393–8400.

6. Lehninger AL, Nelson DL, & Cox MM (1993) Principles of biochemistry.

## References

1. O’Nuallain B, et al. (2006) Kinetics and Thermodynamics of Amyloid Assembly Using a High-Performance Liquid Chromatography–Based Sedimentation Assay. Methods in Enzymology, eds Indu K & Ronald W (Academic Press), Vol Volume 413, pp 34–74.

2. Kuipers BJH & Gruppen H (2007) Prediction of Molar Extinction Coefficients of Proteins and Peptides Using UV Absorption of the Constituent Amino Acids at 214 nm To Enable Quantitative Reverse Phase High-Performance Liquid Chromatography-Mass Spectrometry Analysis. Journal of Agricultural and Food Chemistry 55(14):5445–5451.

3. O’Nuallain B, Shivaprasad S, Kheterpal I, & Wetzel R (2005) Thermodynamics of Aβ(1-40) Amyloid Fibril Elongation†. Biochemistry 44(38):12709–12718.

4. Chen S, Berthelier V, Yang W, & Wetzel R (2001) Polyglutamine aggregation behavior in vitro supports a recruitment mechanism of cytotoxicity. Journal of molecular biology 311(1):173–182.

5. Kar K, et al. (2013) β-Hairpin-Mediated Nucleation of Polyglutamine Amyloid Formation. Journal of Molecular Biology 425(7):1183–1197.

6. Chou PY & Fasman GD (1974) Conformational parameters for amino acids in helical, β-sheet, and random coil regions calculated from proteins. Biochemistry 13(2):211–222.

7. Kim CA & Berg JM (1993) Thermodynamic [beta] -sheet propensities measured using a zinc-finger host peptide. Nature 362(6417):267–270.

8. Minor DL & Kim PS (1994) Measurement of the [beta]-sheet-forming propensities of amino acids. Nature 367(6464):660–663.

9. Hortschansky P, Christopeit T, Schroeckh V, & Fändrich M (2005) Thermodynamic analysis of the aggregation propensity of oxidized Alzheimer’s β-amyloid variants. Protein Sci 14(11):2915–2918.

10. Blondelle SE, Forood B, Houghten RA, & Pérez-Payá E (1997) Polyalanine-based peptides as models for self-associated β-pleated-sheet complexes. Biochemistry 36(27):8393–8400.

11. Robertson AL, Bate MA, Androulakis SG, Bottomley SP, & Buckle AM (2010) PolyQ: a database describing the sequence and domain context of polyglutamine repeats in proteins. Nucleic acids research:gkq1100.

12. Finn RD, et al. (2013) Pfam: the protein families database. Nucleic acids research:gkt1223.

